# Characterizing the effect of impeller design in plant cell fermentations using CFD modeling

**DOI:** 10.1101/2024.11.05.622012

**Authors:** Vidya Muthulakshmi Manickavasagam, Kameswararao Anupindi, Nirav Bhatt, Smita Srivastava

**Affiliations:** Department of Biotechnology, Bhupat & Jyoti Mehta School of Biosciences, Indian Institute of Technology Madras, Chennai-600 036, Tamil Nadu, India; Department of Mechanical Engineering Indian Institute of Technology Madras, Chennai-600 036, Tamil Nadu, India; Department of Data Science and Artificial Intelligence, Wadhwani School of Data Science and Artificial Intelligence, Indian Institute of Technology Madras, Chennai-600 036, Tamil Nadu, India; Robert Bosch Centre for Data Science and Artificial Intelligence, Indian Institute of Technology Madras, Chennai-600 036, Tamil Nadu, India

**Keywords:** Computational fluid dynamics, Plant cell suspension cultures, Stirred bioreactors, Rushton impeller, marine impeller, setric impeller

## Abstract

Cultivation of plant cell cultures in conventional bioreactors designed for microbial cells often results in the decrease of biomass productivity as compared to that in shake flasks, presumably due to the imbalance between the mass transfer requirements and compromise with cell viability. Hit and trial methods are usually adapted to achieve high biomass productivity in the bioreactor. In this study, a rational approach has been developed to choose a suitable impeller for *Viola odorata* cell suspension culture using computational fluid dynamics (CFD). A two-phase CFD model was employed to characterize the non-Newtonian fluid dynamics in a stirred tank reactor using different impeller designs, a setric, Rushton and marine impeller. The simulations were performed adopting Euler-Euler approach for the two-phase flow and dispersed *κ* − ε turbulence model. The numerical model was validated with good agreement with experimental determination of volumetric mass transfer coefficient. The impact of impeller design was then investigated on critical process parameters like mixing, oxygen mass transfer and shear. The developed CFD model demonstrated that setric impeller is a suitable choice for *V. odorata* cell cultivation offering low-shear environment at equivalent velocity magnitudes at reactor bottom with higher cell-lift capabilities which is preferable in high cell-density plant cell cultivations.

## Introduction

Plant cell cultures have been established to be a potential source for consistent and continuous production of diverse high value low volume phytochemicals for a wide variety of industrial applications^1,2^. However, mass cultivation of plant cell cultures in bioreactors for maximum biomass productivity is still considered challenging, presumably due to the relatively large size of the cell, and its ability to form aggregates. This results in wall adhesion, insufficient mixing and cell settling in the bioreactors. High-density plant cell cultivations may exhibit non-Newtonian rheology which could warrant higher agitation rates which is limited by plant cell shear sensitivity^3^. With the aim to maximize plant cell biomass productivity, initially shake flask cultivation is generally done for cell line development where the suitable media and environmental parameters such as initial pH and temperature are optimized. The cultivation is then translated to the bioreactor for optimizing the reactor operating parameters for maximizing biomass productivity overcoming nutrient limitations. But unlike in microbial fermentations, in plant cell cultivations it is observed that the biomass productivities generally reduces when we translate from shake flask to conventional reactor systems^4-7^. In order to overcome growth limitations and obtain atleast comparable biomass productivities in the bioreactor to that of shake flask, hit and trial methods are employed. It includes the use of modified impellers in stirred bioreactors^8-11^ or the use of modified reactor geometrical configurations such as orbitally shaken bioreactors^12^ and wave bioreactors^13^. With the numerous impeller designs and reactor configurations to choose from, and the relatively slow growing nature of plant cell cultures, with a doubling time of approximately 2 to 3 days and a batch time ranging from 3 to 4 weeks, any hit and trial method for a given plant species can be highly time consuming, laborious, incurring huge costs. Therefore, there is a need for a systematic and predictive approach for choosing suitable reactor design and operating conditions *in silico*. This can be achieved using computational fluid dynamics (CFD) which is an excellent tool where the governing equations of fluid flow are solved to provide quantitative and qualitative insights into mixing, mass transfer and the prevailing shear environment in bioreactors^14^. This can facilitate *in silico* design of bioreactor suitable for plant cell cultivation that can offer good mixing and mass transfer at low shear^15^. In spite of the development of few novel plant cell bioreactor designs, and use of pneumatically driven airlift and bubble column bioreactors for plant cell cultivation, stirred tank bioreactors often are the preferred choice as they have well-established scale up principles and facilitate excellent mixing and overcoming mass transfer limitations during scale up of high-density viscous plant cell suspension cultures^3^. Previously, the well-known commonly used Rushton impeller that exhibits radial flow has been used for successful lab scale cultivation of *Sphaeralcea angustifolia*^16^ and *Helianthus annuus*^17^. However, Treat et al. ^18^ observed that during cultivation of soybean and slash pine using stirred bioreactor with Rushton impeller, though biomass productivity was as expected, the cell viability was reduced by 85%. Further, *Panax ginseng* was successfully grown in 50 L bioreactor with marine impeller at an agitation rate of 420 rpm^19^. A specially designed setric impeller has been adopted to successfully cultivate *Podophyllum hexandrum*^20^, *Azadirachta indica*^8^ and *Linum album*^21^ in lab scale bioreactors. With impeller choice being species specific, dependent on cell morphology, cell density and shear sensitivity, it can be beneficial to identify a suitable impeller by characterizing the hydrodynamic environment associated with it in terms of mixing, mass transfer and shear during plant cell cultivation. Hence, this study aims to characterize and quantify these key parameters in the three impellers (Rushton, marine and setric impeller) using CFD modeling for rationally choosing a suitable impeller for plant cell cultivation. Previously, Liu et al^22^. had addressed the effect of shear on plant cells using CFD and assumed that the bubble induced shear forces on plant cell is negligible due to its large size, justifying their use of a single-phase simulation. This study aims to evaluate effect of impellers in stirred bioreactors in terms of hydrodynamics and oxygen mass transfer. Consequently, the system was modeled comprising of two phases: air and the plant cell suspension which accounts for the non-Newtonian nature of plant cells. Further, the developed model was applied onto a lab scale bioreactor as the computational domain with the cultivation of *Viola odorata* plant cell suspension cultures in bioreactors^23^ was considered as a model system for the said CFD analysis. This is the first study characterizing hydrodynamics in bioreactors using CFD for a plant cell suspension with non-Newtonian characteristics to the best of the knowledge of the authors.

## Materials and methods

### Development of the CFD model for stirred tank bioreactor

The section first presents the governing equations that were developed to obtain the fluid flow in a stirred bioreactor for plant cell cultures. Further, the description of the computational domain and boundary conditions applied are discussed. The discretization schemes and the numerical methods used for solving the governing equations are also presented.

### Modeling governing equations for the two-phase fluid flow

The fluid properties of the plant cell suspension was determined experimentally as described in section “Experimental determination of viscosity and average aggregate size of *V. odorata* cell suspension”. It was defined as the primary phase, while the sparged air was defined as the dispersed secondary phase with properties defined at temperature 26.6 °C as this temperature was previously established to be the optimum temperature for maximum biomass productivity in *V. odorata* cell cultures^23^. The two phases were assumed to be incompressible and considered as independently interacting fluid continuum and modeled using the two-phase Eulerian model. The average size of the air bubbles was determined by the Sauter mean diameter correlation (*d*_*b*_) modeled by Kazakis et al. for a porous sparger^24^ as given by:

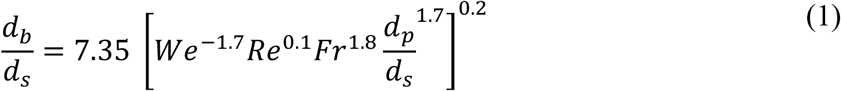

where *We, Re* and *Fr* corresponds to the Weber, Reynolds and Froud numbers respectively, *d*_*s*_ and *d*_*p*_ correspond to diameter and pore size of the sparger, respectively.

The volume (*V*_*q*_) occupied by phase q, in the system is given by Equation (2), as follows:

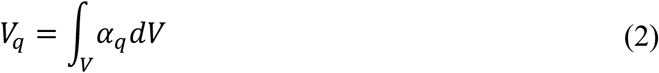

such that *α*_*q*_, the volume fraction of phase q in a single computational element is a continuous function in space and time and hence the sum of volume fractions of all defined phases in a single computational cell is equal to 1 as defined by phasic volume fraction given by Equation (3), as follows:

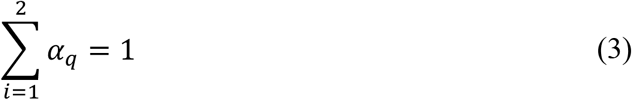

The effective density of phase q in every element is defined as 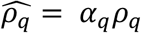 where *ρ*_*q*_ is the physical density of phase q. The Reynolds number of the overall system was calculated to be 3000 as per following equation:

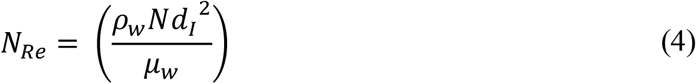

where *ρ*_*w*_ refers to the density of water, *N* refers to the agitation rate, *d*_*I*_ refers to the diameter of the impeller and *μ*_*w*_ refers to the dynamic viscosity of water. The impeller Reynolds number for a bioreactor is generally defined such that greater than 10000 can be generally considered to be turbulent while less than 4, laminar for most cases^25^. Though flow in this system of consideration may fall under transitional regime, modeling the flow as fully turbulent resulted in closer alignment with the experimental data. Hence, the Reynolds-averaged Navier-Stokes (RANS) equations were solved for the two-phase system in this study. A similar approach has also been adopted by Sarkar et al^26^. where they defined critical Reynolds number as 4200 for their system. The multiple reference frame method was adopted to solve the governing equations for the bioreactor^27^. Briefly, the continuity and momentum equations were solved in the stationary reference frame for the bulk region and in the rotating reference frame for the region surrounding the impeller, wherein the instantaneous velocity and pressure values are replaced by the sum of time averaged mean and fluctuating components. The continuity equation was solved for the secondary phase, air, as given by:

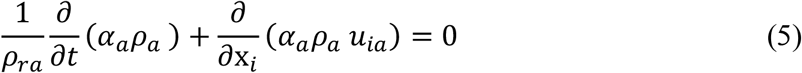

where *ρ*_*ra*_ is the volume averaged density, *ρ*_*a*_ refers to density of air, *α*_*a*_ refers to volume fraction of air and *u*_*a*_ refers to velocity of air. The solution of Equation (5) for *α*_*a*_ was used to calculate volume fraction of *V. odorata* (*α*_*ν*_) from Equation (3).

The momentum equation was solved for both phases individually and is defined as given by:

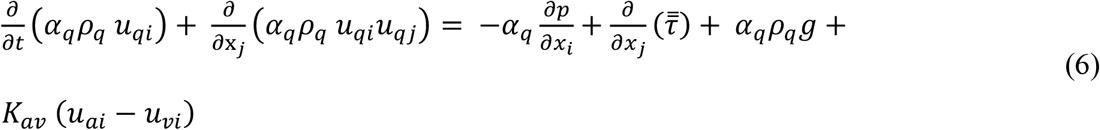

where *α*_*q*_, *u*_*qi*_ and *ρ*_*q*_ are the volume fraction, velocity and density of phase q. p is the pressure force shared between phases, 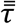 is the stress-strain tensor modeled described as:

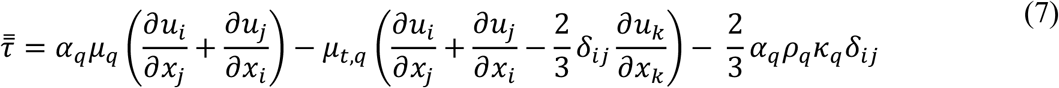

where the turbulent viscosity (*μ*_*t*_) is given by

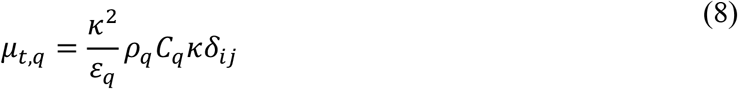

Here, *ε*_*q*_ is the turbulence dissipation rate and *C*_*μ*_ is a constant defined as 0.09. *δ*_*ij*_ is the Kronecker delta function defined as 0 when i ≠ j and 1 when i = j.

*u*_*ai*_ and *u*_*νi*_ in Equation (6) refer to velocities of air and *V. odorata* suspension respectively. *g* corresponds to the gravitational force modeled as 9.81 ms^-2^ in the negative y direction. *K*_*aν*_ refers to the interphase momentum exchange coefficient model between the two phases which tends to 0 when the primary phase is absent in a computational cell and is modeled as

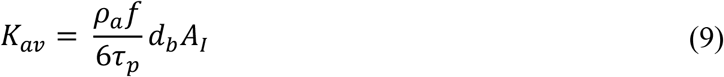

where *A*_*I*_ denotes the interfacial area density, *f* corresponds to the drag function, *d*_*b*_ refers to the air bubble diameter while *τ*_*p*_ models the particulate relaxation time. The interfacial area density for the dispersed phase is modeled as the ratio of surface to volume for a spherical bubble multiplied by the volume fraction of air which is given by

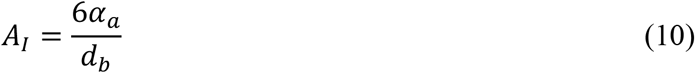

The drag function *f* was modeled by the commonly suitable Schiller Naumann model for most fluid-fluid pairs^27^ which is given by,

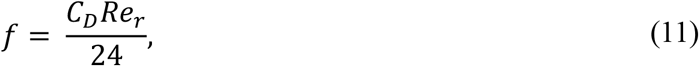

where

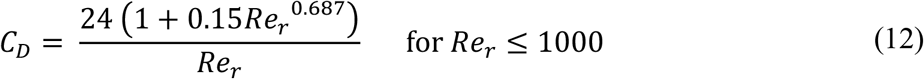

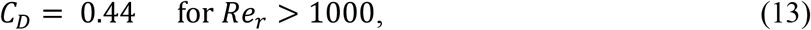

Where *Re*_*r*_, the relative Reynolds number between the primary and secondary phases was calculated by relative velocities of primary and secondary phases and is given by

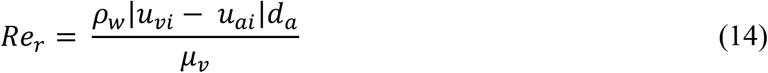

where *μ*_*ν*_ is the viscosity of primary phase. Here, | | refers to absolute value of the difference of relative velocity magnitudes.

The particulate relaxation time (*τ*_*p*_) is modeled as

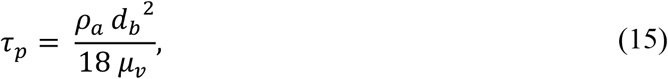

To close the RANS equations, the *κ* – ε turbulence model was employed in this study. The transport equations for turbulent kinetic energy, *κ*, and the turbulent energy dissipation, *ε* for primary phase is given by:

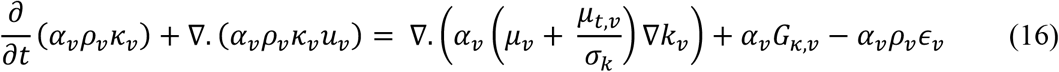

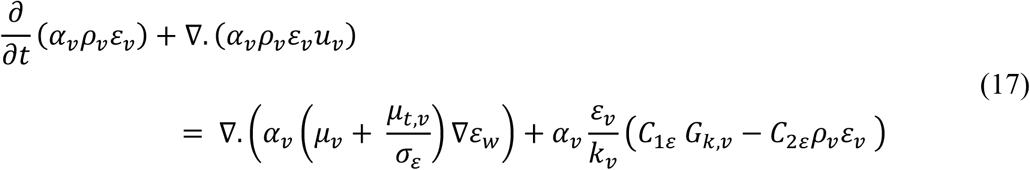

Here, *G*_*k,w*_ refers to the production of turbulence kinetic energy due to mean velocity gradients. Since the volume fraction of the secondary phase is very low compared to the primary phase, air was modeled using the dispersed *κ* – ε turbulence model^27^ in this study. The parameters for the dispersed phase were calculated by assuming homogenous turbulence using the Tchen theory of dispersion of discrete particles^28^. *C*_1*ε*_ and *C*_2*ε*_ are defined as constant values 1.44 and 1.92 respectively, as generally modeled. Further, to predict the velocity gradient at the boundary layer without the use of very fine meshes (with increased computational time), the near wall treatment was modeled using standard wall functions which has been found to be suitable for most flows^27^.

### Details of the bioreactor geometry and computational domain

The geometry of the stirred tank bioreactor was created using ANSYS SpaceClaim 2019R2 (ANSYS, Inc, United States) based on the measurements of a 3.1 L dished bottom glass bioreactor (Applikon Biotechnology B.V, Netherlands) as shown in Figure 1(a). Dished bottom bioreactors are generally recommended for plant cells for better cell lifting capabilities^28^. The tank has a diameter of 130 mm and a total height of 250 mm. The liquid working volume was 2.4 L which corresponded to a tank height of 200 mm. This was taken to be the maximum height of the computational domain in the CFD model to reduce computational effort. A single impeller was employed in this study and the setric, Rushton and marine impeller with design as shown in Figure 1(b)-(d), were placed with an impeller clearance length of 65 mm for adequate mixing of the cell suspension in independent simulations. The setric impeller was custom made at Indian Institute of Technology Madras, and has four blades each inclined at an angle of 60° with a diameter of 45 mm, blade length 49 mm and width of 12 mm^29^. Rushton and marine impellers were standard impellers provided by Applikon Biotechnology B.V, Netherlands. A porous sparger has been used for sparging air in this study as it has the capacity to generate small bubbles with increased oxygen exchange area at low gas rates suitable for plant cell cultivation^28^. The sparger (Applikon Biotechnology B.V, Netherlands) with diameter of 6 mm (Figure 1(e)) and a pore size of 15 μm was placed 4 mm below the shaft of the impeller. Three baffles of length 139 mm and width 12 mm were placed in the bioreactor to prevent vortex formation. The fluid domain was only considered for the simulation and the solid components were removed from the fluid domain as void spaces using the split body option in SpaceClaim software. In order to model the rotation of the impeller using multiple reference frames, a cylindrical fluid domain 1.25 times the impeller diameter called the rotating domain was created enclosing the impeller, as shown in Figure 1 (a). The region outside the rotating domain was considered stationary. To achieve a conformal meshing between the rotating and stationary domain, topology was shared between the two bodies. The computational domain was then discretized into finite volumes using ANSYS Meshing 2019 R2 (ANSYS, Inc, United States). The geometry was meshed using unstructured tetrahedral mesh with refinement near the impeller region with 0.68 million computational cells which had an average skewness and orthogonality of 0.25 and 0.75 respectively and a maximum aspect ratio of 4. The grid size was chosen after a grid sensitivity study was performed for seven grid sizes ranging from 0.2 to 2.5 million to ascertain that the obtained results are independent of the mesh (supplementary Fig. S1). In addition, the grid convergence index was calculated to confirm the asymptotic convergence of the chosen grids (supplementary Table TS1). This mesh was used for further simulations and the computational domain used is shown in Figure 1 (b).

**Figure 1:**
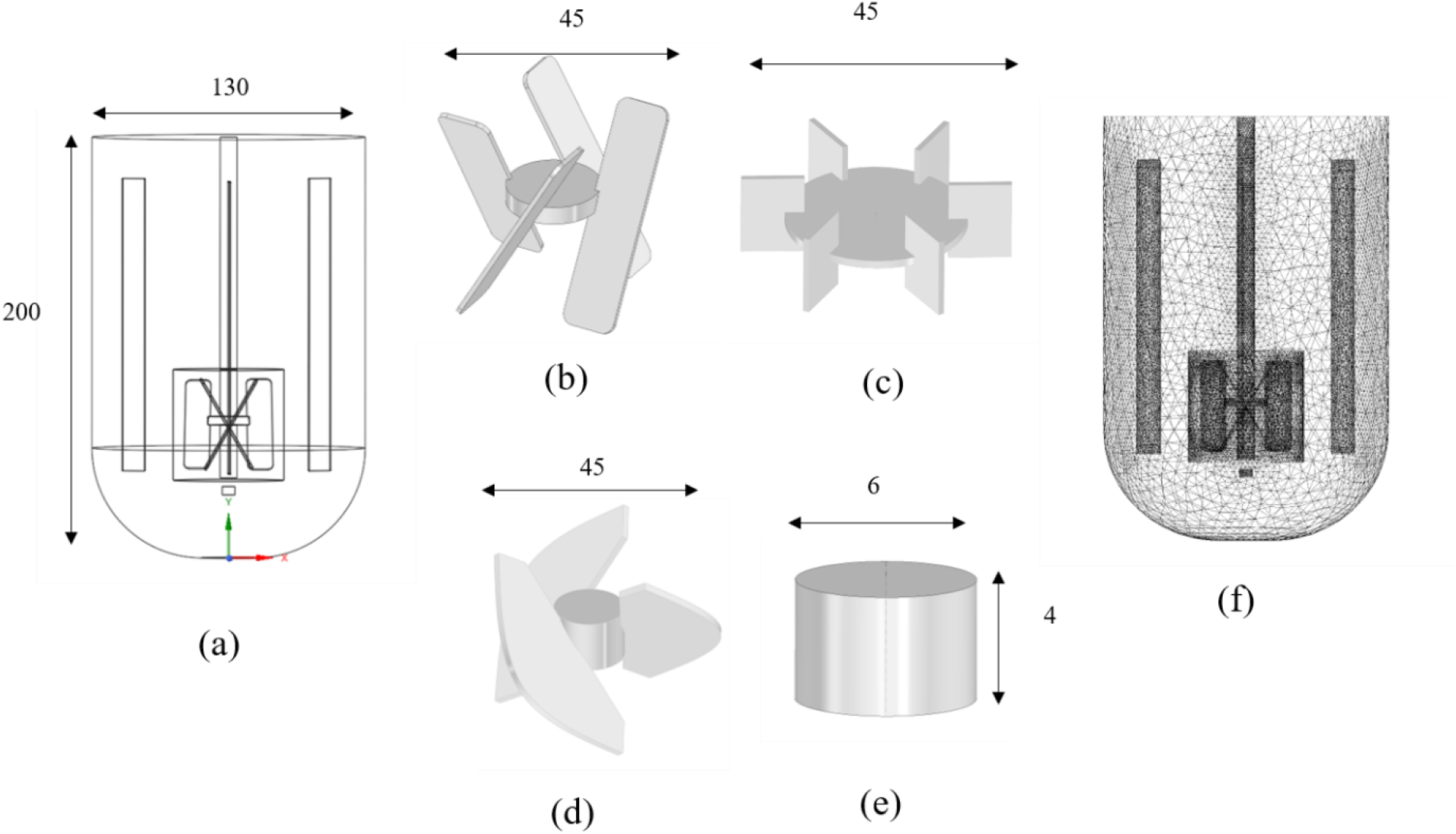
Schematics of the geometry of the stirred tank bioreactor (3 L) with relevant measurements (a) Schematic of the simulated bioreactor geometry in the XY plane (b) setric impeller (c) Rushton impeller (d) marine type impeller (e) Meshed computational domain of the simulated bioreactor geometry in the XY plane (All dimensions are in mm)

### Definition of boundary conditions and numerical methods for solving the governing equations

The governing equations were constrained by boundary conditions at the computational domain limits. The liquid level (maximum limit of the computational domain) was considered to be a degassing boundary wherein only air is allowed to escape from the liquid. The walls of the bioreactor vessel, shaft, impeller and the sparger wall were considered to be under no-slip condition for both air and the plant cell suspension. The inlet region of sparger was considered to be the input surface for air bubbles at an aeration rate of 0.2 vvm (volume of air per unit volume of culture medium per minute) and the impeller cell zone was set to an agitation rate of 85 rpm. This condition was adopted from cultivation conditions of *V. odorata* cell cultures^23^ to compare the hydrodynamics in all three impellers. Since the temperature of the system is maintained constant for plant cell growth, and being an incompressible flow, the pressure-based solver was used to solve the governing equations. Here, the pressure corrector equation, phase coupled-SIMPLE algorithm^27^ was used to correct the velocity to overcome the constraint of solving Equation (5) simultaneously with Equation (6). The gradient term was spatially discretized using least squares cell-based algorithm. The pressure equation was discretized to second order and the other governing equations were discretized to first order accuracy. The temporal discretization was performed with implicit integration with first order accuracy. Initially, a single phase (water only) simulation was carried out. The single-phase steady state results were used to initialize air-water simulation which was further used to initialize the air-*V. odorata* simulation to overcome convergence difficulties. The time step was taken to be 1.5 ms with maximum 20 iterations per time step. The governing equations of the model with appropriate boundary conditions were solved numerically using the commercially available code, ANSYS Fluent 2019R2 (ANSYS, Inc, United States) in a high performance computing environment. Convergence was defined to be attained when the residuals reached an order of 10^−5^ for all equations and a constant volume averaged value of k_L_a, energy dissipation rate and liquid velocity were achieved through subsequent time steps.

### Modeling of volumetric liquid phase oxygen mass transfer coefficient

Volumetric mass transfer coefficient of oxygen (k_L_a) is a parameter generally used to characterize the mass transfer of oxygen from the air to the liquid phase in stirred tank bioreactors. It is calculated as the product of the liquid mass transfer coefficient *k*_*L*_ and the interfacial area *a*. In this study, the Higbie’s penetration model^30^ was used to determine *k*_*L*_, which was also successfully used for modeling *k*_*L*_ by Sarkar et al. ^26^ and Amer et al. ^31^ for air-water fluid pairs given by

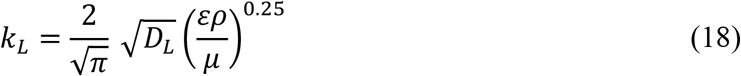

where *D*_*L*_is the molecular diffusivity of oxygen in liquid. For a non-Newtonian fluid, Equation (18) was modified by Kawase and Moo^32^ as given by

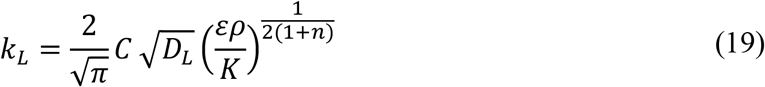

Here C was taken as 0.15^33^. The interfacial area available for mass transfer between phases was calculated as given in Equation (10). Here, K and n refers to the consistency and flow behavior index respectively. These equations were incorporated in Fluent using a custom field function.

### Modeling of shear stress acting on plant cell cultures

Plant cells are exposed to shear stress based on its relative size to the Kolmogorov length scale. The Kolmogorov length (*η*_*k*_) is given by (*ν*^3^/*ε*)^0.25^ where *ν* and *ε* stand for kinematic viscosity and liquid energy dissipation rate respectively. If the cell size is smaller than *η*_*k*_, the shear stress to which the cell is exposed to is controlled by the hydrodynamics within the eddy and given by^22^:

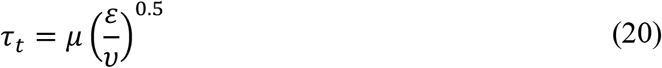

Unlike microbial and mammalian cells, plant cells tend to form aggregates and if the aggregate size is larger than *η*_*k*_, potentially ranging from 0.1 – 2 mm in diameter^22^, the cells are subjected to shear due to velocity differences across the diameter of the cell aggregate. This shear stress can be calculated as given in the following equation^22^

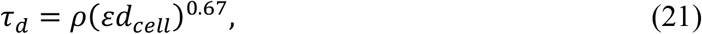

where *d*_*cell*_ refers to the diameter of the cell aggregate. These equations were incorporated in Fluent using a custom field function.

### Experimental determination of volumetric mass transfer coefficient (k_L_a) for model validation

In this study, *k*_*L*_*a* for setric impeller was experimentally determined using dynamic gassing in method^34^. Briefly, the bioreactor (3.1 L) with setric impeller was autoclaved with 2.4 L working volume of water and connected to the DO sensor for polarization. After cooling, the bioreactor was aerated to constant oxygen concentration at 125 rpm and 2 vvm. One point calibration at 100% was performed after the saturation was sufficiently reached. The bioreactor was then stripped off oxygen completely by purging nitrogen gas. Once the dissolved oxygen concentration reached less than 2%, filtered compressed air was sparged in the reactor at the simulated operating conditions until steady state saturation was reached. The measurements of dissolved oxygen were recorded as a function of time using a polographic oxygen sensor (Applikon Biotechnology B.V, Netherlands). To calculate the rate of oxygen transfer from gas to liquid, the mass balance for the dissolved oxygen in the bioreactor is given by:

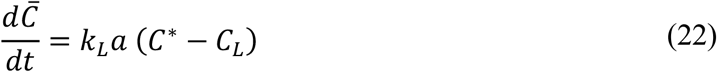

Here, *C*_*L*_ refers to the dissolved oxygen concentration and *C*^*^ refers to the saturation concentration of oxygen in water calculated based on Henry’s Law^25^. Integrating Equation (22), from t=0 to saturation time, we have

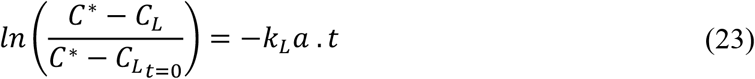

Here t is time in seconds and 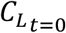 is given by the oxygen concentration at the start of the experiment (less than 2%). *k*_*L*_*a* was estimated from the slope by plotting 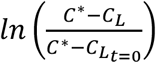 as a function of t from Equation (23) using linear regression in Microsoft office excel 2019.

### Experimental determination of viscosity and average aggregate size of V. odorata cell suspension

*V. odorata* VOP-4 callus was previously established by Narayani et al.^35^ and was maintained by periodic subculturing. The cell suspension culture was developed as described by Babu et al^23^. Briefly, the maintained callus (6 gDW/L) was suspended in 50 mL woody plant medium (Himedia Laboratories) with 3% (w/v) sucrose and 3 mg L^-1^ of 2,4-dichlorophenoxy acetic acid at an initial pH of 5.8. The cells were grown in an orbital shaker in conical flasks for 14 days at 23°C with a photoperiod of 16/8 h light/dark cycle. Further, the cells were filtered using a Buchner funnel to obtain a synchronous fine cell suspension which was used as inoculum for cultivation of *V. odorata* in bioreactors^23^. In order to bring the CFD model closer to reality, the rheological parameters of the cell suspension were characterised experimentally. *V. odorata* cell suspension was subjected to varying shear rates 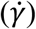 from 0.0095 -1000 s^-1^ to determine the corresponding shear stress (*τ*) using the rheometer AntonPaar MCR502 at 25°C. The flow behavior (n) and consistency index (K) were determined correlated by the Ostwald de Waele equation given by

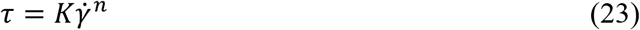

The experiment was repeated at n=5 and the data is presented as mean ± standard error. Further, in order to obtain the average cell aggregate size of *V. odorata* in suspension, 20 *μ*L of the suspension was pipetted using cut tips onto a glass slide and was observed under an Olympus IX83 inverted fluorescence microscope. The images were then processed using the software ImageJ^36^ to determine the range and average cell aggregate size.

## Results and Discussion

### CFD model convergence and validation

The CFD model for the air-water system was first verified and validated as the steady state results of this simulation was used to initiate the air-*V. odorata* cell suspension system. It was ascertained that the residuals of all the governing equations decreased below an order of 10^−5^, and the monitored parameters, including the volume averaged velocity, energy dissipation rate and k_L_a achieved steady state (supplementary Fig. S2). Additionally, the mass conservation of the inlet and the outlet air flow rate was confirmed at the sparger inlet and degassing boundary. Since the multiple reference frame method has been used as the boundary condition for the CFD simulation, rather than actual movement of the impeller, the impeller tip speed from the CFD model was verified with theoretical tip speed for all three impellers. Theoretical tip speed of the impellers were calculated to be 0.2 ms^-1^ from the equation *u*_*tip*_ = *π*. *N* d_*I*_, and the maximum liquid velocity from the CFD simulation at the impeller was found to be 0.212, 0.212, 0.195 ms^-1^ for setric, marine and Rushton impeller, respectively which is considerably close to the theoretical value with an error of less than 6%. The CFD model was then verified with the literature reported ungassed power number for Rushton impeller as the correlation between impeller Reynolds number and the corresponding power number is well-established for the air-water system^25^. The Reynolds number of 3000 calculated from Equation (4) corresponded to a power number of 4.5 and the power number estimated from the present CFD simulation was found to be 3.17. The error of 30% could be justified by the use of 4 baffles in the experimental study in contrast to 3 used in this study. Further, the CFD predicted *k*_*L*_*a* in the air-water simulation was validated with experimentally determined *k*_*L*_*a* at an agitation rate of 85 rpm and aeration rate of 0.2 vvm for setric impeller. A less than 4% error in k_L_a was found between the experimental and numerical data for the air-water system as shown in Table 1, confirming the validity of the model for further analysis. To validate the air-non-Newtonian system, the *k*_*L*_*a* determined from CFD simulation was validated using the literature reported experimental *k*_*L*_*a* at an agitation rate of 125 rpm and aeration rate of 0.2 vvm for setric impeller^8^. It was observed that CFD predicted k_L_a had an error of 22% compared to the experimental data for the air-plant cell suspension system as shown in Table 1. This could be attributed to a variation in the rheological properties of the plant cell suspension between the experimental and CFD study.

**Table 1:**
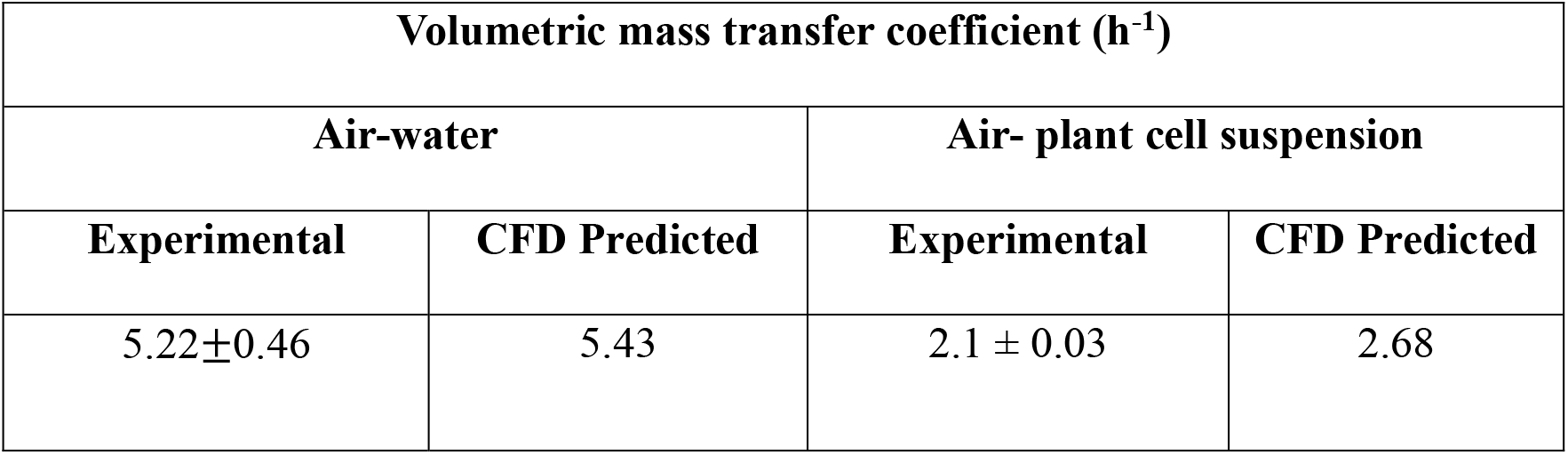
Comparison of volumetric mass transfer coefficient (k_L_a) for setric impeller determined experimentally and predicted from CFD simulations for model validation.

| Volumetric mass transfer coefficient ($\text{h}^{-1}$) | | | |
| --- | --- | --- | --- |
| Air-water |  | Air- plant cell suspension |  |
| Experimental | CFD Predicted | Experimental | CFD Predicted |
| $5.22 \pm 0.46$ | 5.43 | $2.1 \pm 0.03$ | 2.68 |

The validated CFD model was then used for analyzing the impact of using different impellers on the fluid dynamics, oxygen mass transfer and shear in the bioreactor to rationally choose the suitable impeller for the cultivation of the model system *V. odorata* without performing hit and trial experiments.

### Experimental determination of rheological parameters and the range of cell aggregate size for V. odorata cell suspension

The experimentally determined rheological parameters of *V. odorata* from the shear rate-stress data fit to the Ostwald de Waele equation, presented in Table 2 was used to model the liquid in the CFD simulation with properties. With a flow behavior index less than 1, *V. odorata* cell suspension was found to be pseudoplastic in nature. This characteristic is similar to other plant cell species which also exhibit pseudoplastic behaviour^37,38,39^. The non-Newtonian behavior of plant cells is generally attributed to cell elongation in suspension and the ability of plant cells to grow in high cell densities due to its comparatively lower oxygen demand^40^. In addition, the cells tend to aggregate and the average size and range estimated from image analysis from microscopic observations is tabulated in Table 2.

**Table 2:**
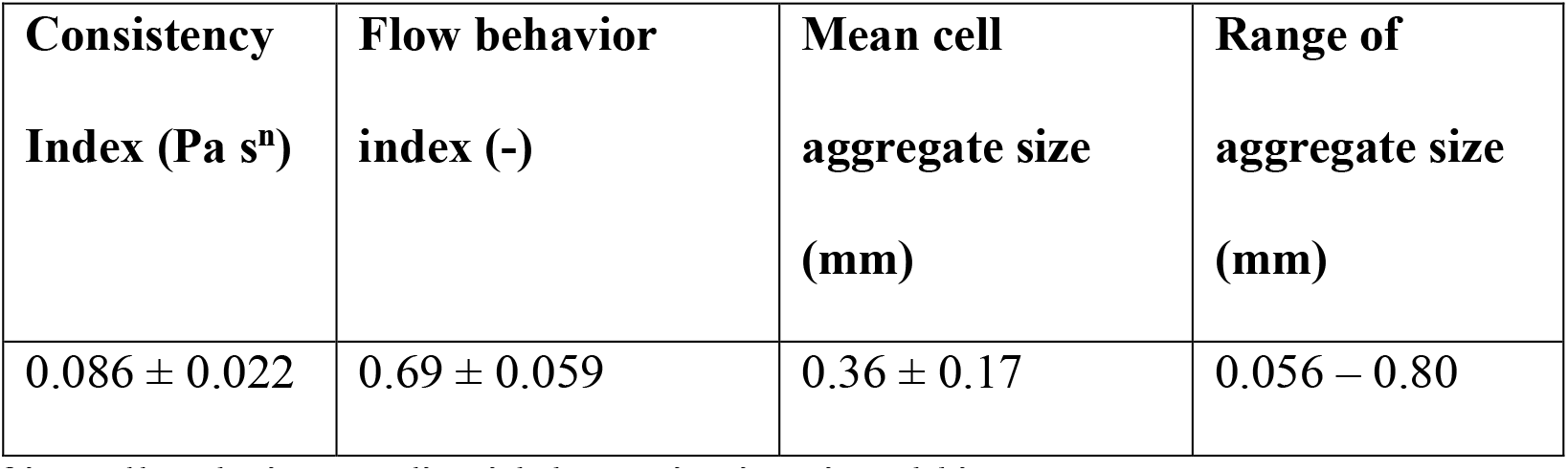
Rheological properties, cell aggregate size and range of *V. odorata* cell suspension cultures on day 0 (inoculum) used in the CFD simulation to model the non-Newtonian behavior of the fluid and determine shear due to cell aggregate size.

| **Consistency**<br>**Index (Pa s<sup>n</sup>)** | **Flow behavior**<br>**index (-)** | **Mean cell**<br>**aggregate size**<br>**(mm)** | **Range of**<br>**aggregate size**<br>**(mm)** |
| --- | --- | --- | --- |
| 0.086 ± 0.022 | 0.69 ± 0.059 | 0.36 ± 0.17 | 0.056 – 0.80 |

### Effect of impeller design on liquid dynamics in stirred bioreactor

With viscous plant cell suspension cultures, ineffective mixing can lead to development of dead zones which can hamper effective nutrient availability which is undesirable^41^. Thus, for plant cells, the mixing efficiency of the impeller is assessed by the dead zone volume within the bioreactor in this study. The fluid flow patterns obtained by the effect of three different impellers in the stirred bioreactor in the air-plant cell suspension systems can be observed in the contour/velocity plots in Figure 2. A dead zone in the bioreactor can be defined as the regions where the liquid velocity is less than 5% of the impeller tip speed^42^. Based on this definition, the blue regions in Figure 2, with velocity magnitudes below 0.011 ms^-1^ can be interpreted as the dead zones. In all three impeller configurations, it can be observed that the change in rheology from Newtonian (supplementary Fig. S3) to pseudoplastic (Figure 2) in the presence of plant cells has decreased the impeller mixing efficiency leading to increase in dead zones. The fluid in the setric impeller configuration is pushed axially downward forming a recirculation loop towards the impeller region demonstrating that the impeller has predominantly downward pumping axial flow (supplementary Fig. S3). It is interesting to note that the fluid flow pattern shifts from fully axial (in air-water (supplementary Fig. S3)) to a combination of axial and radial (in air-*V. odorata* (Figure 2 (a)) which is in line with previous reports that axial impellers tend to show radial behavior when agitating highly viscous non-Newtonian fluids^43^. The same can be observed by the radial increase in the impeller cavern in the axial marine impeller configuration ((Figure 2 (b)). However, the marine impeller pumps the fluid upward in contrast to the downward pumping setric impeller which is in line with experimental observations of Afedzi et al^44^. The simulations of Rushton impeller demonstrated the expected radial profile of the fluid movement (Figure 2 (c)) and the characteristic formation of trailing vortices behind the flat blade of the impeller (Figure 2 (f))^25^.

**Figure 2:**
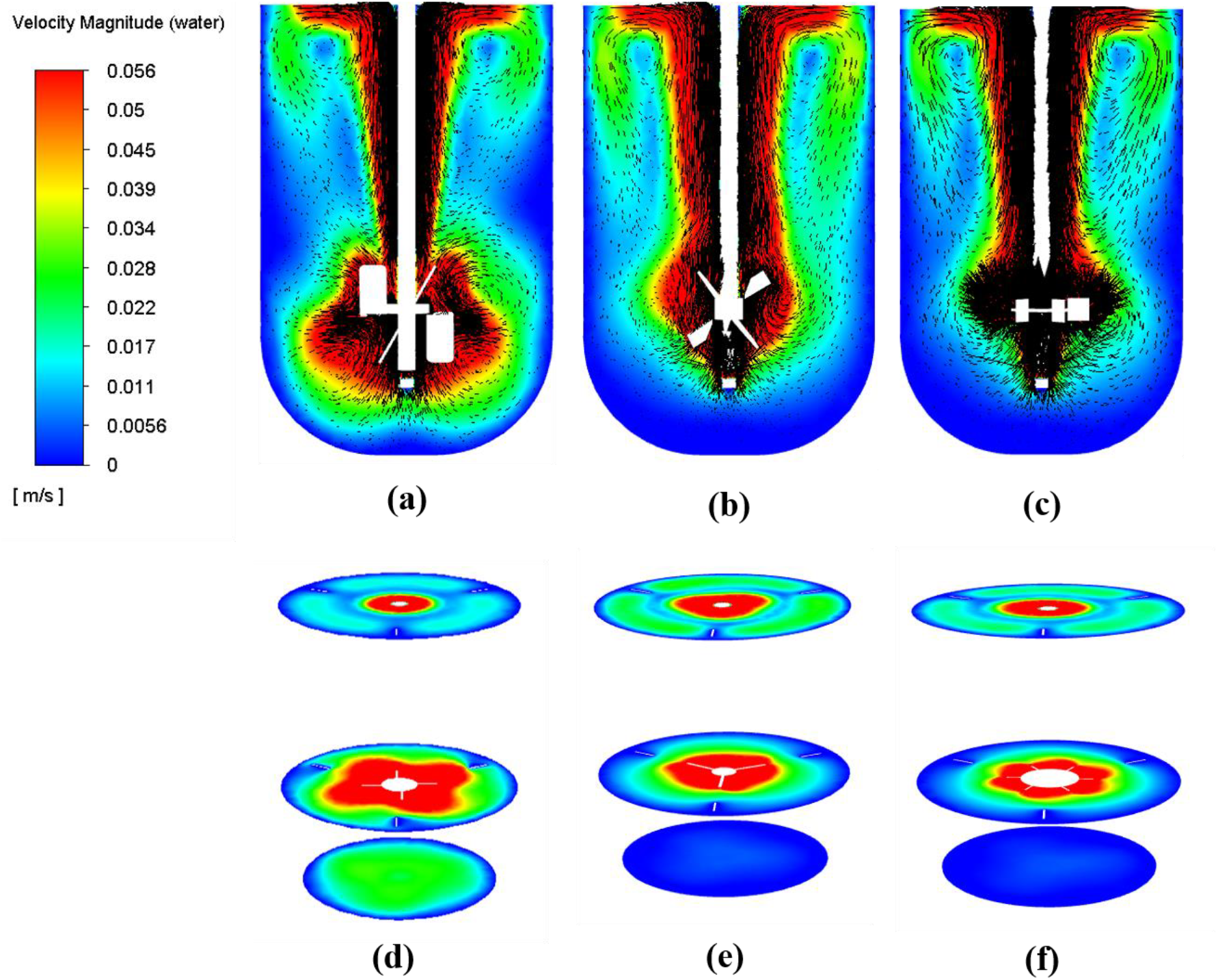
Contours and vector plots of fluid velocity across a vertical plane in an air-plant cell suspension system with different impellers ((a), (b), (c)) and across three different horizontal planes (d), (e) and (f) at agitation rate of 85 rpm and aeration rate of 0.2 vvm., (d) correspond to setric impeller (b), (e) correspond to marine impeller (c), (f) correspond to Rushton impeller (The number of arrows is not an indication of mesh density)

It is critical to prevent plant cell aggregates from settling down with gravity as that could eventually lead to necrosis due to insufficient nutrients and oxygen during cultivation in bioreactors^3,45^. In this regard, it is critical to have good fluid movement in the regions below the impeller devoid of dead zones. In the presence of plant cells, it can be observed that setric impeller is able to facilitate good velocity magnitudes up to 0.02 ms^-1^ (10% of the tip speed) in the bottom plane, 20 mm from the base of the reactor (Figure 2 (d)). In contrast, at this agitation and aeration rate, both marine and Rushton impeller facilitate only 0.0056 ms^-1^ (less than 3% of the impeller tip speed) at the bottom plane (Figure 2 (e) and (f)) leading to negligible fluid movement. At the plane above the impeller (150 mm from the base of the bioreactor) the liquid is discharged after encountering the liquid surface, on to the walls and recirculates in two loops in each side of the vessel axially from top to bottom. This second recirculation loop that moves the fluid upward, primarily directed due to the sparged air is almost similar in all three impellers as the aeration rate is constant among the three cases. At the top plane, therefore all three impellers facilitate an average velocity magnitude of 0.03 ms^-1^ (15-17% of impeller tip speed) (Figure 2 (d), (e) and (f)) indicating the potential for good fluid movement for the plant cells. At the plane across the impeller, (65 mm from the base of the bioreactor) it is evident that the setric impeller exhibits a higher average velocity magnitude compared to the Rushton and marine impellers (Figure 2 (d), (e) and (f)), with fluid movement sweeping well across until the bioreactor walls. In contrast, both Rushton and marine exhibit localized mixing only in the vicinity of the impeller. This could lead to adherence of plant cells on the walls due to lack of movement, in addition to cell settling which is highly unfavorable^46^.

To further substantiate this observation across the entire volume of the reactor, in addition to analyzing at three planes, the dead zone volume was calculated by dividing the bioreactor into four compartments as illustrated in Figure 3 (a). Figure 3 (b) demonstrates that at the bottom setric impeller facilitated 56% lesser dead zone volume than Rushton and marine. Similarly, below the impeller, setric facilitated 74% and 71% lesser dead zone volume than Rushton and marine respectively. At the top of the reactor, all three impellers offered effective mixing with less than 15% dead zone volume similar to the plane wise analysis. These results indicated that the observations at the three planes are true for the behavior in the corresponding compartments as well.

**Figure 3:**
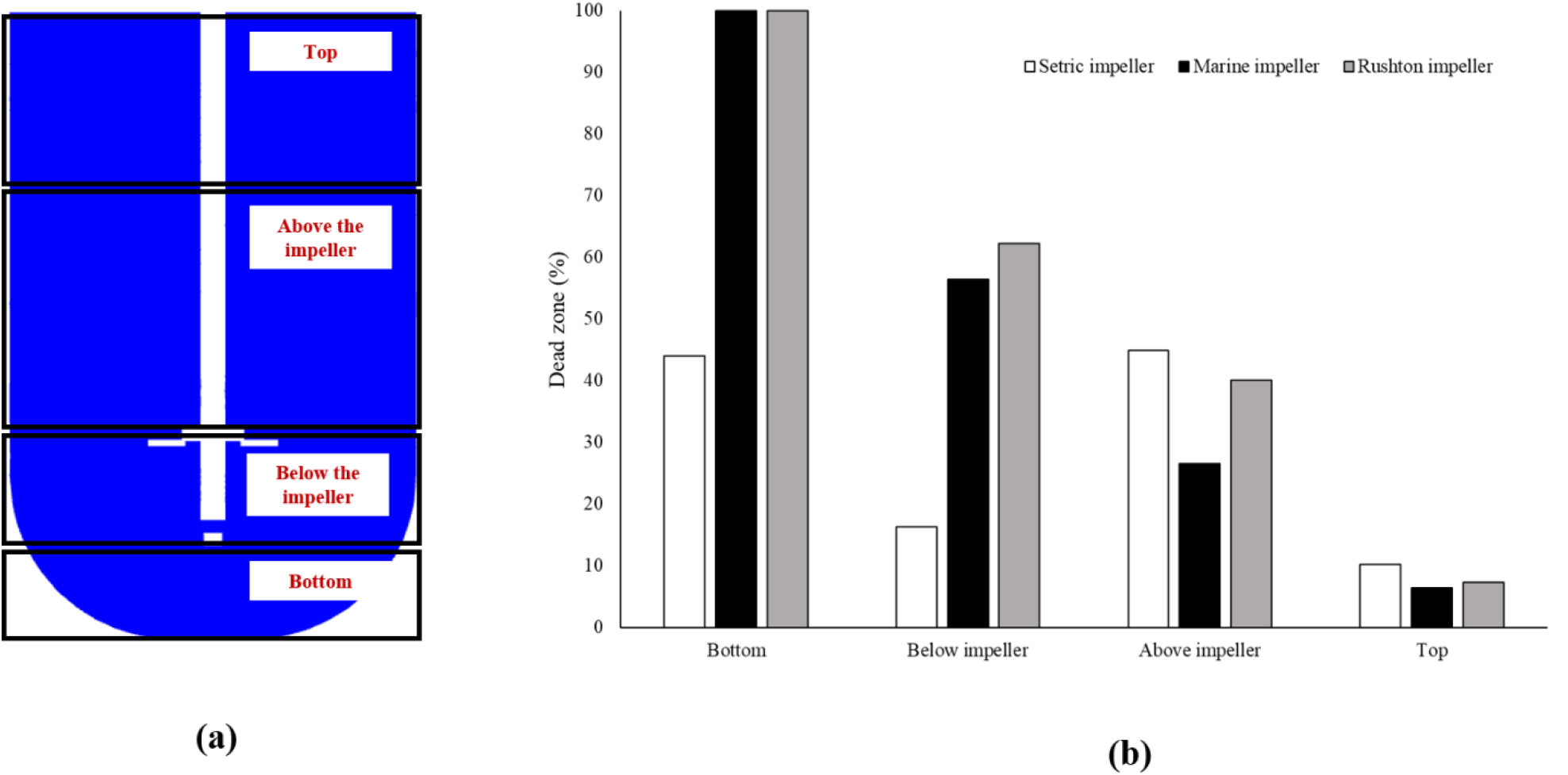
Calculation of dead zone volumes in the 3 L stirred tank at agitation rate of 85 rpm and aeration rate of 0.2 vvm (a) Compartment divisions in the bioreactor for analysing localised dead zone percentages (b) Localised pecentage dead zones in setric, Rushton and marine impeller at the four different compartments

Though both Rushton and marine impellers are able to facilitate good mixing in air-water system throughout the bioreactor, the dead zones have increased substantially in presence of plant cells particularly below the impeller. It is crucial to prevent cell settling and wall growth when designing bioreactors suitable for plant cells, and hence this comparative analysis from the CFD study indicates that the setric impeller configuration facilitates effective mixing in the reactor bottom and sufficient mixing in the top regions, promoting efficient cell movement throughout the reactor and making it a suitable choice for cultivation of *V. odorata* cell suspension culture.

### Effect of impeller design on liquid velocity distribution in stirred bioreactor

To further compare the hydrodynamic patterns among the three impellers in presence of plant cells, the distribution of liquid velocity components on a line at three different spatial locations in the reactor were studied, including one below the impeller (20 mm from base of the bioreactor) (Figure 4(a)), second at the impeller center (65 mm from base of the bioreactor) (Figure 4(b)) and third at the top of the impeller region (150 mm from base of the bioreactor) (Figure 4(c)). The axial velocity, component of fluid parallel to the length of the bioreactor demonstrates how effectively the cells circulate through the bioreactor, crucial for adequate oxygen and nutrient availability. This is particularly important for plant cell aggregates due to their relatively higher settling velocity. In this regard, at the reactor bottom, setric impeller could facilitate a maximum axial velocity of 0.022 ms^-1^ and a radial velocity of 0.024 ms^-1^ which is close to four times higher axial and radial velocities than Rushton and marine impeller (Figure 4 (d) and (e)). It is important to note that the magnitudes of axial and radial velocities exhibited by the setric impeller is almost similar to the average bottom velocity magnitude (0.023 ms^-1^) and setric impeller offers axial liquid movement near the bioreactor walls and radial movement in the bottom center. This gives an insight that the fluid pushed down from the impeller radially moves towards the bioreactor wall before recirculating back to the impeller region. This fluid movement facilitated by setric impeller is clearly favorable for preventing plant cells from settling.

**Figure 4:**
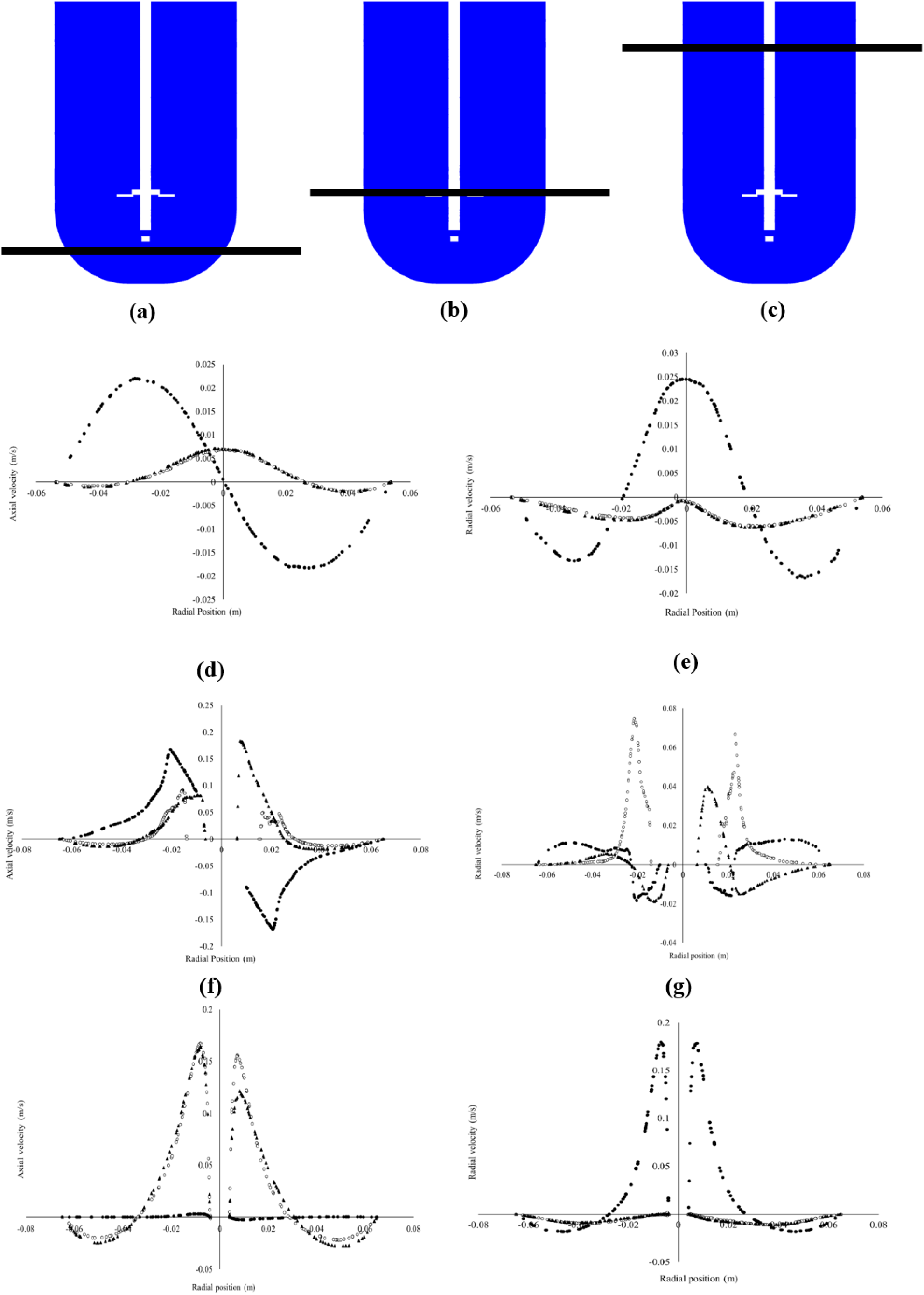
Effect of impeller configuration on axial and radial velocity along radial positions at three different lines in the stirred bioreactor (a) at 20 mm (b) at 65 mm (c) at 150 mm from the base of the bioreactor vessel at agitation rates of 85 rpm and aeration rate of 0.2 vvm (d) and (e) represent the axial and radial velocity respectively at a line located 20 mm upwards to the base of the bioreactor, which represents the bottom zone of the bioreactor; (f) and (g) represent the axial and radial velocity respectively at a line located 65 mm upwards to the base of the bioreactor, which represents the impeller zone of the bioreactor; (h) and (i) represent the axial and radial velocity respectively at a line located 150 mm upwards to the base of the bioreactor, which represents the top zone of the bioreactor (● distribution of axial/radial velocity magnitude (m s^-1^) in setric impeller, ▴ distribution of axial/radial velocity magnitude (m s^-1^) in marine impeller, ○ distribution of axial/radial velocity magnitude (m s^-1^) in Rushton impeller)

The primary behavior of the impeller can be assessed by examining the axial and radial velocities at the impeller center^9^, which can provide insight into the nature of fluid movement in different vessel geometries, scales, or when multiple impellers are utilized.

Figure 4 (f) show the axial velocity distributions for the three different impellers at the impeller region. The model indicated that the setric impeller offered a maximum of 1.8 times higher axial velocity magnitude (0.167 ms^-1^) than Rushton and similar magnitudes to marine impeller in the upward direction. On the other hand, in the downward direction setric impeller exhibits a maximum axial velocity magnitude of 0.168 ms^-1^ (equivalent to the upwards axial velocity magnitude), which is 14 times higher than the maximum axial velocity magnitude of Rushton and 10 times higher than that of the marine impeller. It is evident that at the impeller region, setric impeller moves the fluid in both up and downward direction facilitating good mixing in contrast to Rushton and marine impellers which moves the fluid only towards the top. On the other hand, the radial velocity magnitude in Rushton impeller (0.074 ms^-1^) was upto 2 times and 4 times higher than marine and setric respectively which is as expected as it is well known that Rushton impeller exhibits radial behavior predominantly (Figure 4 (g)). But it is interesting to note that the magnitudes of radial velocity in Rushton impeller is 2.2 times lower than that of axial velocity (0.167 ms^-1^) in setric impeller at the impeller region which indicates stronger fluid movement in the impeller region with setric configuration. The radial velocity in marine impeller exhibits similar fluid movement to setric impeller throughout the monitored region, with two times higher magnitudes near the impeller.

This confirms that at the impeller region, the primary motion of the fluid can be considered axial for setric impeller. It is noteworthy that the marine impeller exhibits comparable axial velocity magnitudes to setric solely in the upward direction, whereas the setric impeller facilitates fluid movement axially in both upward and downward directions. It is reported that axial impellers are preferred over radial impellers for suspending plant cells in general^28^ and non-Newtonian fluids^43^. The observations from this CFD study substantiate that radial impellers cannot recirculate fluid effectively at the bottom, and consequently, axial impellers are preferred.

Furthermore, it can be indicated that downward-pumping axial impellers are more suitable, as it can be seen that upward-pumping marine impellers cannot recirculate fluid sufficiently towards the bottom as well.

Analyzing the velocity component distributions at a line in the top region (Y=150 mm), it was observed that the maximum axial velocities were 50 times higher in the upward direction for the Rushton and marine impeller than the setric impeller configuration (0.0033 ms^-1^) which had negligible axial velocity ((Figure 4 (h)). This could be attributed to the liquid movement majorly directed towards the surface by the sparged air without radial movement in the Rushton and marine impeller configurations. This observation indicates that the higher velocity magnitudes in the top region, as depicted in Figure 3 (b) for Rushton and marine impellers, are primarily axial. Conversely, the setric impeller facilitated up to a maximum of 0.179 ms^-1^ at the top which was 16 times higher than Rushton and marine impellers (Figure 4 (i)). These results indicate that although the average velocity magnitudes are lower at the reactor top for the setric impeller, as observed in Figure 2 (b), the radial velocities are higher than Rushton and marine impellers, potentially providing effective recirculation for the plant cells as well as oxygen distribution.

Notably, in all three locations examined, the axial component of the fluid movement is predominantly towards the top for Rushton and marine impellers, whereas the setric impeller facilitates a balance between the upward and downward movement, indicating effective mixing. These studies indicated that setric impeller is predominantly a downward pumping axial impeller which is found to be better than Rushton and marine impellers in facilitating cell lift. This is especially critical at relatively lower agitation rates which are typically employed for plant cell cultivations to maintain cell viability^41^.

### Effect of impeller design on the volumetric oxygen mass transfer coefficient and shear stress in stirred bioreactor

Though the model indicated that setric impeller facilitates better mixing at the reactor bottom making it a suitable choice for plant cell suspension cultures, it is important to additionally consider and compare oxygen mass transfer and shear environment in the bioreactor as they are critical parameters for cultivation of plant cells in bioreactors. The average k_L_a and shear stress in the bioreactor in presence of the three impellers was compared keeping the average liquid velocity magnitude in the reactor bottom constant as this parameter cannot be compromised.

The average bottom velocity in setric impeller at 85 rpm was 0.02 m/s which was taken as the benchmark. The agitation rate at which bottom velocity magnitude in Rushton and marine impellers were similar (less than 15% error) to setric at 85 rpm was identified by running the bioreactor simulations at increasing rpm from 100-300 rpm. It was observed that at 330 rpm, the average bottom velocity in Rushton impeller was 0.0214 and at 300 rpm, the average bottom velocity in marine impeller was 0.0204 as tabulated in Table 3. At this agitation rate and constant aeration of 0.2 vvm, the oxygen mass transfer and the shear stress were compared in the following sections.

**Table 3:** CFD estimated Kolmogorov length scale ranges for the three different impellers at similar average velocity magnitudes at reactor bottom tabulated along with corresponding average velocity and energy dissipation rates.

| Impeller<br>configuration | Agitation<br>rate (rpm) | Average<br>velocity<br>magnitude<br>at reactor<br>bottom (m/s) | Average<br>velocity<br>magnitude<br>(m/s) | Energy<br>dissipation<br>rate (m<sup>2</sup>/s<sup>3</sup>) | Kolmogorov length (mm) |  |  |
| --- | --- | --- | --- | --- | --- | --- | --- |
|  |  |  |  |  | Min | max | Mean |
| Setric | 85 | 0.0232 | 0.0277 | 0.0166 | 0.085 | 9.62 | 2.99 |
| Marine | 300 | 0.0204 | 0.0579 | 0.0427 | 0.063 | 5.6 | 2.07 |
| Rushton | 330 | 0.0214 | 0.0492 | 0.0476 | 0.046 | 5.9 | 2.24 |

### Effect of impeller design on the volumetric oxygen mass transfer coefficient at constant average velocity magnitude at reactor bottom

Oxygen mass transfer is one of the crucial limiting factors in aerated fermentations. Plant cells require relatively less oxygen than microbial cultures as they are slow growing and have low specific oxygen demand^3^. The rate of oxygen mass transfer from gas bubbles to the liquid is dependent on concentration gradient of oxygen (difference between the maximum soluble oxygen concentration in the liquid and the actual oxygen concentration in the bulk liquid) and k_L_a^25^. Among the two, k_L_a is the parameter which is controllable. It is dependent on impeller type, aeration and agitation rate and therefore it is important to compare the same between the three impellers. In this study, the model indicated that at the similar average velocity magnitude at the bottom, it was observed that Rushton impeller demonstrated 1.8 times better oxygen mass transfer than setric (1.37 h^-1^) and marine impeller (1.32 h^-1^) as seen in Figure 5(a). This can be understood by the ability of radial Rushton impellers to disperse bubbles efficiently, leading to better interfacial area and better k_L_a^25^. The marine impeller had a similar magnitude of k_L_a to setric impeller even at 300 rpm due to a similar interfacial area in spite of increase in the turbulence dissipation rate. The model hence indicates that in terms of oxygen mass transfer, Rushton impeller is the most suitable among the three impellers. However, there is generally a tradeoff between better oxygen mass transfer and low shear in bioreactor design. Since both parameters are functions of energy dissipation rates, when the impeller dissipates more energy, with high shear at the impeller, it leads to better bubble dispersion and better k_L_a. But since plant cells are shear sensitive and have much lower oxygen demand than microbial cells, it is more crucial to identify the impeller which offers lower shear environment than better oxygen mass transfer for maximum plant cell biomass productivity.

**Figure 5:**
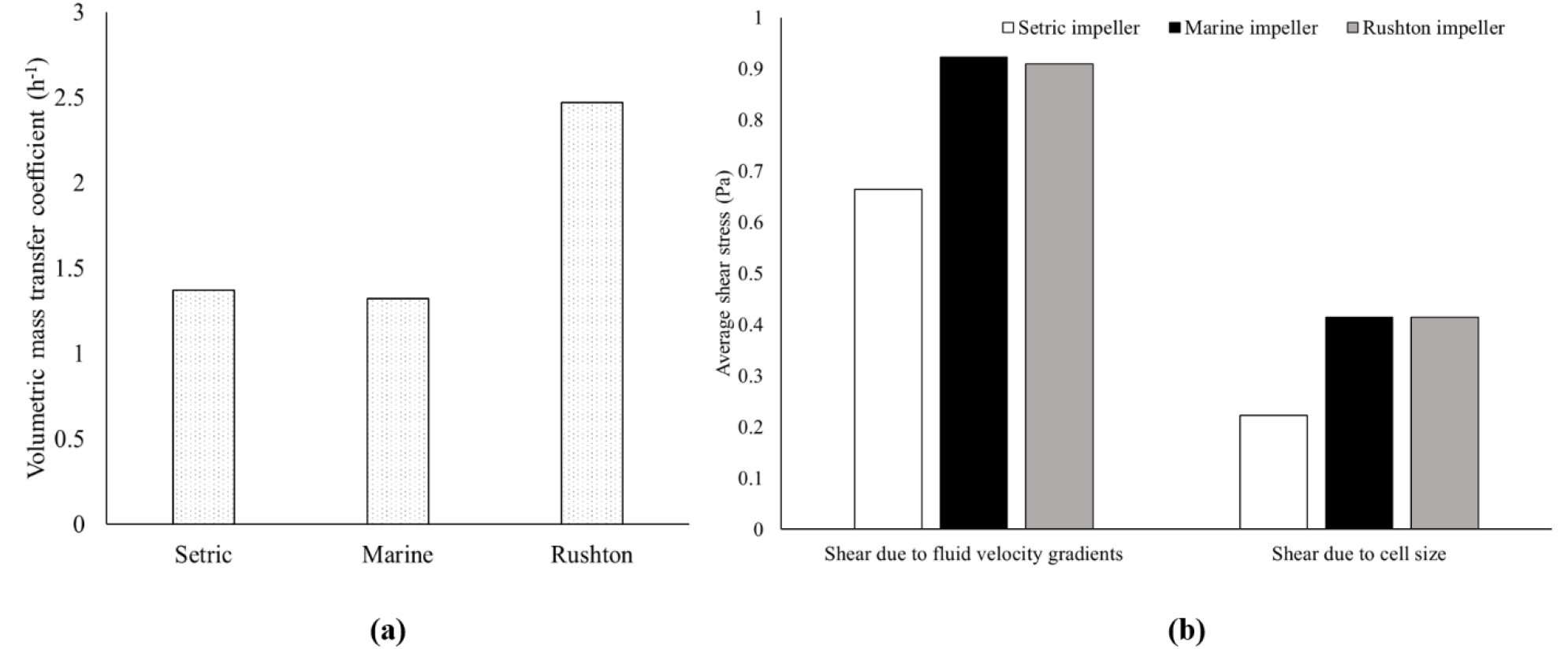
Effect of impeller design on the volumetric oxygen mass transfer coefficient (k_L_a) and shear stress in stirred bioreactor at similar magnitudes of average velocity magnitude at the bottom region of the bioreactor (a) Effect of impeller design on average k_L_a determined from CFD simulations calculated based on Higbie’s penetration model (b) Effect of impeller design on average shear stress due to fluid velocity gradients and shear stress due to cell size calculated from CFD simulations

### Effect of impeller design on the shear environment in the bioreactor at constant average velocity magnitude at reactor bottom

The two approaches generally adopted to determine shear environment in the bioreactor: shear due to velocity gradients (***τ***_***t***_) and shear due to aggregate size (***τ***_***d***_)^22^ has been employed in this study. The shear environment among the three impellers was evaluated at comparable average velocity magnitude at the bottom of the reactor, similar to the approach used for oxygen mass transfer assessment. Figure 5 (b) illustrates the average magnitudes of ***τ***_***t***_ and ***τ***_d_. It can be observed that ***τ***_***t***_, in setric impeller (0.66 Pa) is 1.4 times lower than that of Rushton and marine impellers. On the other hand, ***τ***_d_ exhibited by setric impeller (0.22 Pa) is close to 1.9 times lower than Rushton and marine impeller. The Kolmogorov length scale is the determining factor in whether a cell/cell cluster experiences ***τ***_***t***_ or ***τ***_d_ in the flow field, with the estimated range varying between 0.046 and 9.6 mm in this system. The corresponding ranges for each impeller configuration has been tabulated in Table 3. In this study, it was estimated that the diameter of *V. odorata* cell aggregates ranges between 0.056 – 0.802 μm during inoculation (Table 2) and therefore cell aggregates in this system can experience both ***τ***_***t***_ or ***τ***_d_ depending on their size.

However, comparing the mean cell aggregate size of *V. odorata* (0.36 mm) with the mean Kolmogorov length scales (Table 3) in this system, it can be observed that cell aggregates could most likely experience ***τ***_***t***_ in the flow field. Therefore, it can be indicated from the model that choosing an impeller with lower ***τ***_***t***_ is critical for *V. odorata* and therefore setric impeller can be a favorable choice over the other two impellers.

## Conclusion

There is a need for rational design of bioreactors for plant cell bioreactors rather than adopting existing conventional designs through hit and trial approaches, saving time and resources. In this study, the hydrodynamics, mixing and mass transfer in stirred bioreactors using setric impeller has been characterized using CFD and compared with commonly used axial marine and radial Rushton impeller at relatively lower agitation and aeration rates generally used for plant cell cultivations with a fluid exhibiting plant cell characteristics for the first time. The CFD analysis indicated that setric impeller exhibited considerable downward pumping of fluid and facilitated higher velocity magnitudes below the impeller region than Rushton and marine impeller. These characteristics of setric impeller can prevent cell settling, which is commonly observed in plant cell cultivation as plant cells tend to grow as aggregates. Further, the model indicated that at similar velocity magnitudes at the reactor bottom, setric impeller exhibits lower average shear stress. Based on the fact that lower shear environment is preferred to enhanced oxygen mass transfer for plant cells, the CFD analysis was able to indicate that setric impeller facilitates low shear environment with good mixing, making it a better choice vis a vis the commonly used Rushton turbines and marine impellers for shear sensitive plant cell cultivations. This investigation was substantiated by the choice of setric impeller in a previous experimental study for mixing high density *V. odorata* cell cultivation in a stirred bioreactor where setric impeller was able to facilitate shake flask reproducible biomass productivity in the bioreactor at the same operating conditions^29^. It is also important to note that the agitation rate may need to be varied to achieve the same mixing conditions at low shear as the apparent viscosity and the cell aggregate size can increase during the log phase during cultivation. Thus, this developed model can be further extended for rational optimization of the operating conditions during the cultivation period and to improve biomass productivity for scale up of plant cell cultivation from lab to pilot scale.

## Acknowledgements

The authors acknowledge the use of the computing resources provided by HPCE at Indian Institute of technology (IIT) Madras for running simulations. Authors would like to thank Dr. Abhijit P. Deshpande, Professor, Department of Chemical Engineering, Indian Institute of Technology Madras and Ms. Puchalapalli Saveri for their support in measurement of rheological parameters of the *Viola odorata* cell suspension culture. This work was supported by Hyclone Life Sciences Solutions India Private Limited (Cytiva) (Project Number: CR22230114BTHLSS008458) and L&T Technology Services (Project Number: CR21221810BTLNTE008458) under Corporate Social Responsibility (CSR). Author VM has received Prime Minister’s Research Fellowship (PMRF) from the Ministry of Education, Government of India.

## Author contributions

VM: Methodology, Investigation, Data curation, Visualization, Writing – Original Draft. KA: Conceptualization, Formal analysis, Validation, Writing – Review and editing NB Supervision, Conceptualization, Formal analysis, Validation, Writing – Review and editing. SS: Supervision, Funding acquisition, Resources, Conceptualization, Formal analysis, Validation, Writing – Review and editing.

All authors read and approved the final manuscript

## Data availability

All data supporting the finding of this study are available in the paper and its supplementary information section.

## Additional information

