## Supplementary information for "Characterizing the effect of impeller design in plant cell fermentations using CFD modeling"

A grid sensitivity study was performed to ensure that the obtained results are independent of the chosen mesh size. In this regard, three different meshes (N1=2.5, N2=0.68, N3=0.2 million cells) were chosen and volume averaged energy dissipation rate, mass transfer coefficient and the velocity magnitude in setric impellers were monitored over time for 7.5 seconds to evaluate the grid quality as shown in Figure S1.

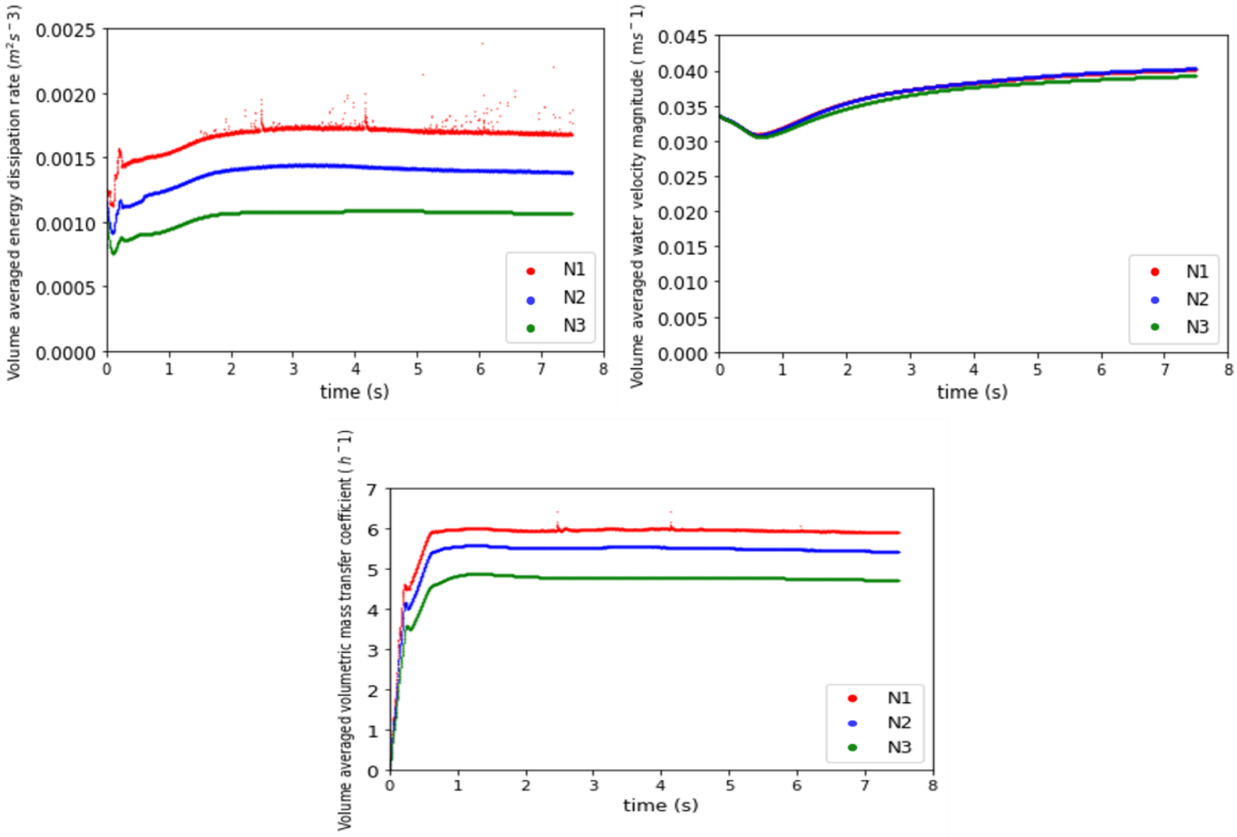


Figure S1: Grid independence study performed for the bioreactor with setric impeller at an agitation rate of 85 rpm and aeration rate of 0.2 vvm to ensure the obtained results are independent of mesh size monitoring three different hydrodynamic parameters with time including (a) Volume average energy dissipation rates at different mesh densities (b) Volume average velocity magnitudes at different mesh densities (c) Volume average volumetric mass transfer coefficients at different mesh densities

In addition, a grid convergence index (GCI) study was conducted using the method reported by Roache. In this study, a small value of GCI indicates that the computational solution falls within the asymptotic range. Briefly, GCI is evaluated by:

$$GCI = \frac{F_S|e|}{r^p - 1} \quad \text{(ES1)}$$

where $F_S$ is the factor of safety which taken as 3 for three or more grids. The error (e) between the parameters considered is calculated between the consecutive grids by:

$$|e| = \left|\frac{f_2 - f_1}{f_1}\right| \quad \text{(ES2)}$$

Here $f$ can be the chosen parameter for the grid sensitivity study. Further the mesh refinement ratio r was calculated as $\frac{N_{i-1}}{N_i}^{1/3}$ and N1 > N2 > N3 where N refers to number of grids in a mesh. The order of convergence (p) was calculated as given:

$$p = ln\left(\frac{f_3 - f_2}{f_{2-}f_1}\right) \quad \text{(ES3)}$$

The calculation is tabulated for volume averaged energy dissipation rate, mass transfer coefficient and the velocity magnitude in setric impellers in Table TS1.

Table TS1: Details of different parameters considered in grid-convergence index study and their calculated values in order to ascertain the asymptotic convergence of the chosen grids

| | **N1 (2.6M)** | **N2 (0.68M)** | **N3 (0.20M)** | | | | | |
|---|---|---|---|---|---|---|---|---|
| **Volume average** | **f1** | **f2** | **f3** | **e12** | **e23** | **p** | **GCI 12** | **GCI 23** |
| **Energy dissipation** | 0.00167 | 0.00138 | 0.00106 | 0.174 | 0.23008 | 0.204 | 2.36 | 3.12 |
| **$k_La$** | 0.00163 | 0.00150 | 0.00131 | 0.079 | 0.13078 | 0.947 | 0.198 | 0.32 |
| **Velocity** | 0.0401 | 0.0402 | 0.0393 | 0.0035 | 0.02417 | 4.468 | 0.00075 | 0.00514 |

The volume average energy dissipation rate, mass transfer coefficient and the velocity magnitude were monitored to evaluate convergence of the CFD model. As shown in Figure S2, the simulations were carried out until a significantly constant value was achieved. The results indicated that the steady state value was reached after 6 seconds, and hence the simulation was run for 7.5 seconds.

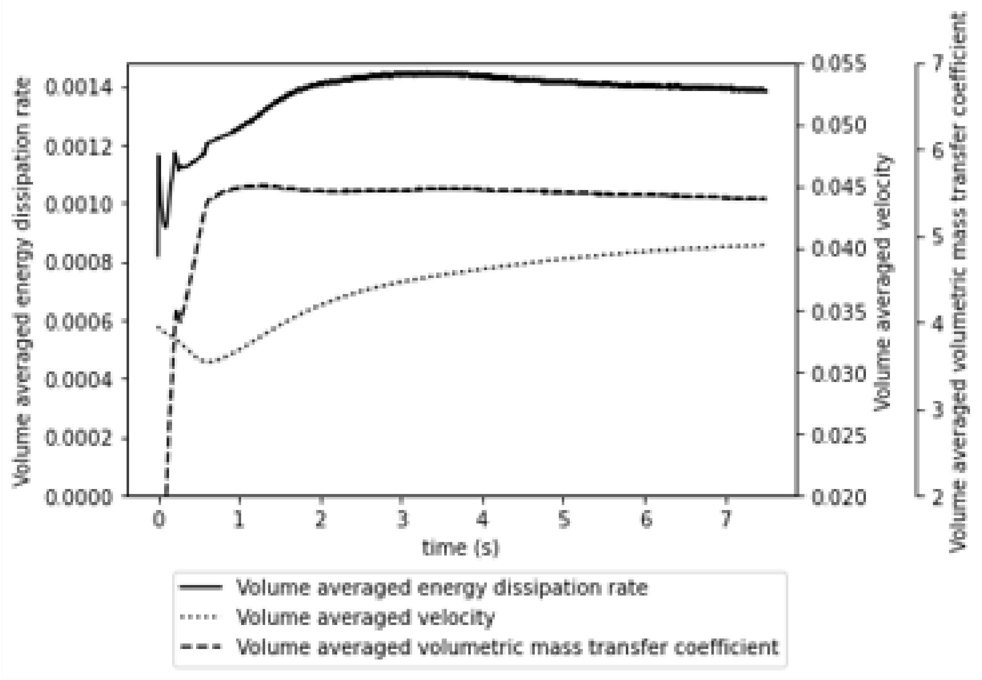


Figure S2: Monitored parameters of volume averaged energy dissipation rate, mass transfer coefficient and the velocity magnitude in stirred bioreactor with setric impeller at an agitation rate of 85 rpm and aeration rate of 0.2 vvm

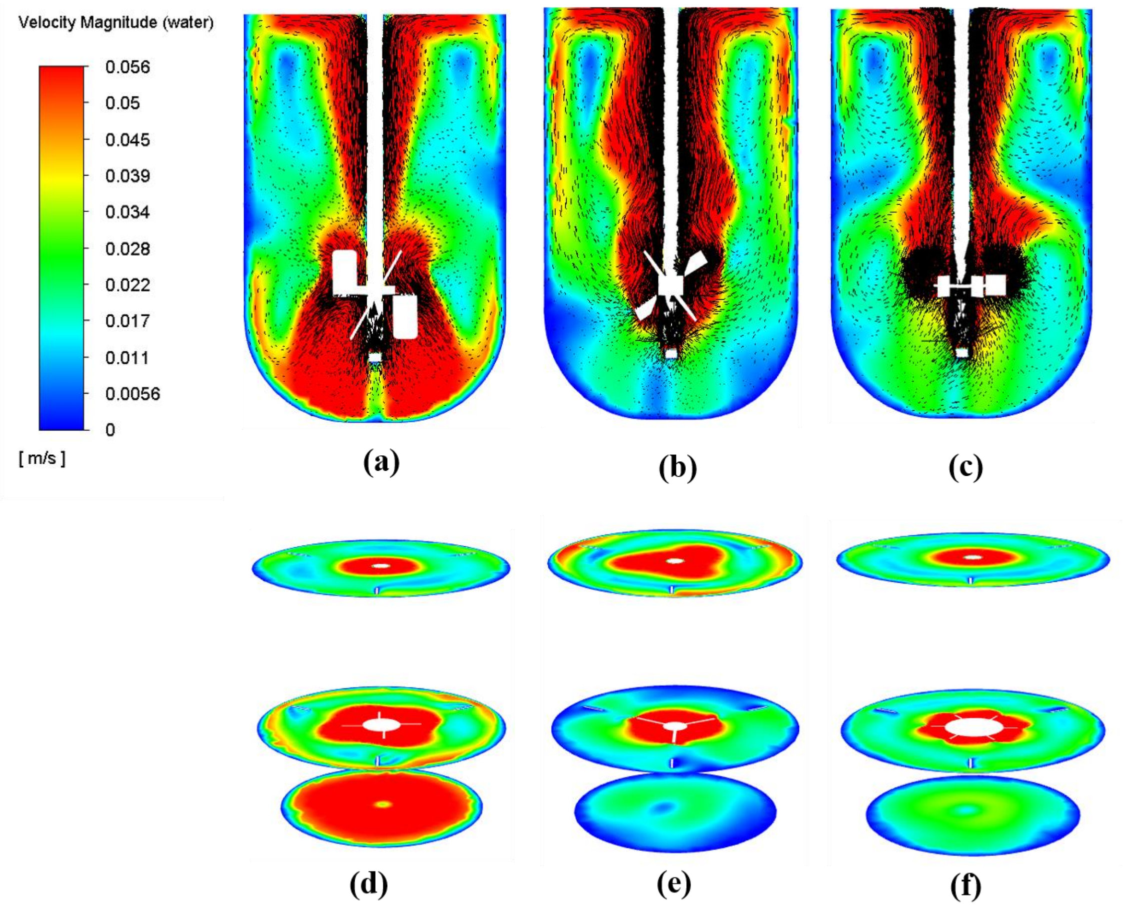


Figure S3: Contours and vector plots of fluid velocity across a vertical plane in an air-water system with different impellers ((a), (b), (c)) and across three different horizontal planes (d), (e) and (f) at agitation rate of 85 rpm and aeration rate of 0.2 vvm.

(a), (d) correspond to setric impeller (b), (e) correspond to marine impeller (c), (f) correspond to rushton impeller (The number of arrows is not an indication of mesh density)
